# *ECSfinder*: Optimized prediction of evolutionarily conserved RNA secondary structures from genome sequences

**DOI:** 10.1101/2024.09.14.612549

**Authors:** Vanda Gaonac’h-Lovejoy, John S. Mattick, Martin Sauvageau, Martin A. Smith

**Affiliations:** Department of Biochemistry and Molecular Medicine, Faculty of Medicine, Université de Montréal, Montreal, QC, H3T 1J4, Canada; CHU Sainte-Justine Research Centre, Montreal, QC, H3T 1C5, Canada; Montreal Clinical Research Institute, Montreal, QC, H2W 1R7, Canada; School of Biotechnology and Biomolecular Sciences, Faculty of Science, UNSW Sydney, NSW 2052 Australia; UNSW RNA Institute, UNSW Sydney, NSW 2052 Australia; Department of Biochemistry and Center for RNA Sciences, McGill University, Montreal, QC, H3G 1Y6, Canada

## Abstract

Accurate prediction of RNA secondary structures is essential for understanding the evolutionary conservation and functional roles of long noncoding RNAs (lncRNAs) across diverse species. In this study, we benchmarked two leading tools for predicting evolutionarily conserved RNA secondary structures (ECSs)—*SISSIz* and *R-scape*— using two distinct experimental frameworks: one focusing on well-characterized mitochondrial RNA structures and the other on experimentally validated Rfam structures embedded within simulated genome alignments. While both tools performed comparably overall, each displayed subtle preferences in detecting ECSs. To address these limitations, we evaluated two interpretable machine learning approaches that integrate the strengths of both methods. By balancing thermodynamic stability features from *RNALalifold* and *SISSIz* with robust covariation metrics from *R-scape*, a random forest classifier significantly outperformed both conventional tools. This classifier was implemented in *ECSfinder*, a new tool that provides a robust, interpretable solution for genome-wide identification of conserved RNA structures, offering valuable insights into lncRNA function and evolutionary conservation. *ECSfinder* is designed for large-scale comparative genomics applications and promises to facilitate the discovery of novel functional RNA elements.

## Introduction

Non-protein coding sequences compose 98% of the human genome, with at least 75% of these sequences being transcribed, mostly into long noncoding RNAs (lncRNAs) that originate intergenically or antisense with respect to protein-coding genes (1–4). In the October 2022 release (v.42) of the GENCODE human gene set annotations, the number of lncRNA genes surpassed that of protein coding genes (5), while over 100, 000 human lncRNAs have been recorded in other databases (6, 7). There are likely many more, given the under sampling of cells at different developmental stages (8) and the high-resolution analyses that have revealed the expression of previously unreported lncRNAs across chromosome 21 (3) as well as from intergenic regions associated with complex traits and diseases (9, 10).

There is considerable evidence that lncRNAs form modular structures (11, 12), supported by the observation that the internal exons of lncRNAs are almost universally alternatively spliced (3). There is also growing evidence that lncRNAs contain conserved structures that interact with specific proteins (4, 13–16). Consequently, there have been a variety of attempts to predict RNA secondary structures from genomic and transcriptomic datasets. However, the uncertain reliability of structure prediction algorithms has fueled debate on the nature and prevalence of evolutionarily conserved structures in lncRNAs (17–20).

RNA secondary structure prediction algorithms have evolved from early thermodynamic models to more recent deep learning approaches. Traditional RNA folding algorithms like *Mfold* (21), *RNAstructure* (22), and *RNAfold* (23) are based on Zuker’s algorithm (24), which uses dynamic programming to identify the most stable RNA folding by minimizing an approximation of the molecule’s global free energy (25, 26). The “energy directed” folding approach is efficient but limited in accuracy due to simplifying assumptions (27) and the inherent nature of global optimization functions, which can ‘overpredict’ RNA structures (28). These limitations, rather than measurement errors, restrict prediction accuracy, which averages only about 67% (29).

The presence of a stable secondary structure in an RNA does not necessarily imply functionality, as random RNA sequences can form complex secondary structures that statistically mirror those associated with known functional elements (30–33). This can be due to several variables, such as base pairing dynamics, transcriptional kinetics, tertiary and quaternary structural interactions, base modifications, and other external factors. This underscores the importance of assessing the evolutionary conservation of secondary structures (ECSs) as a better predictor of functionality. Indeed, identifying the presence of compensatory mutations and base-pair covariations (usually limited to secondary structure via Watson-Crick and G-U ‘wobble’ base pairs) that are consistent with a common higher-order structure across a given evolutionary distance is a hallmark of evolutionary selection for molecular function operating through structural topology.

The identification of conserved RNA structures through multiple sequence alignments has been addressed by several computational methods, including *RNAz* (34), *EvoFold* (35), *CMfinder* (36), *SISSIz* (37, 38) and *R-scape* (20, 39), among others (40–45). Some tools, such as RNAz and *SISSIz*, use an alignment annotated with a consensus secondary structure as input, which is then subjected to probabilistic assessment of evolutionary conservation. The *RNAalifold* algorithm from the Vienna RNA package is commonly used for consensus secondary structure prediction; it ‘folds’ the alignment consensus sequence using a combination of thermodynamic energy and column-specific covariation pseudo-energy scoring metrics (46). A modified version of this algorithm, *RNALalifold*, enables the detection of locally stable consensus RNA structures within a user-defined maximal base pair range (23), avoiding the need to employ fixed-length sliding windows to scan large alignment blocks, which segments sequences and can lead to false negative predictions.

These tools are instrumental in predicting regions within large eukaryotic genomes, such as the human genome, that likely harbor functional RNA structures. However, the sensitivity and specificity of genome-wide predictions using these tools comes with certain reservations. Notably, they have been reported to suffer from poor signal-to-noise ratios and typically operate near the limits of statistical significance (22–24), which complicates the distinction between true evolutionary signal and background noise. Moreover, empirically assessing the performance of these algorithms is challenging due to difficulty in establishing true negative controls, a lack of consensus among different tools, uncertainties in statistical significance calculations, and other methodological issues and assumptions (47).

Building on the success of deep learning in protein structure prediction, similar methods are now being applied to RNA secondary structure prediction (48–51). However, a significant challenge is the scarcity of training data, leading to inflated performance due to overlap between training and testing sets (27, 52, 53). The Critical Assessment of Structure Prediction 15 (CASP15) study has reported that current experimental methods outperform deep learning models for RNA structure prediction (54). Additionally, models with many parameters, such as *ContextFold* (55), often overfit, resulting in low prediction accuracy (56, 57). Thus, the accuracy of predicting secondary structures for RNA sequences dissimilar to the training data is limited unless specific measures are taken to prevent overfitting (27, 49, 53, 58). Moreover, these deep learning models are often challenging to interpret (59).

Here, we describe two experimental frameworks to benchmark RNA secondary structure conservation prediction algorithms in the context of comparative genomic screens. We focus on two leading algorithms that assess statistical significance of proposed consensus RNA secondary structures: *SISSIz* (37) and *R-scape* (20); the former has been used to suggest that there are >4 million evolutionarily constrained RNA structures detectable in multiple genome alignments of 35 mammals (60), whereas the latter finds no statistically significant support for evolutionary conservation of proposed secondary structures of the functionally validated lncRNAs *HOTAIR*, *SRA*, and *Xist* (*20*). We leverage well-annotated ncRNAs in mitochondrial genome alignments of eutherian mammals (61) to compare the accuracy of both tools. We also assess their performance using a benchmarking pipeline engineered to reproduce the empirical conditions of comparative genomic screens via shuffled genomic alignments harboring sequences sampled from experimentally validated Rfam structures (62).

To address the challenges posed by thermodynamic, covariation, and deep learning approaches, we developed *ECSfinder*. This tool employs a random forest (RF) classifier to harmonise the strengths of *SISSIz* and *R-scape* in a highly interpretable manner by integrating biophysical properties and evolutionary data from homologous sequences. By merging these established principles with evolutionary insights, *ECSfinder* aims to offer a robust and transparent solution for RNA secondary structure prediction, addressing the shortcomings of deep learning models and enhancing the predictability and reliability of the process.

## Materials and methods

### Algorithms

#### SISSIz

The *SISSIz* algorithm addresses the issue of high false positive rates in RNA structure prediction by generating dinucleotide-controlled phylogeny-aware shuffled alignments, which preserve base-pair stacking energies. It models site-specific nucleotide interactions based on the branch lengths from a phylogenetic tree inferred from the original alignment. A Z-score is generated by comparing the *RNAalifold* consensus score (a measure of thermodynamic stability and compensatory base-pairs) to a normal background distribution generated from multiple iterations of SISSI (37). The *SISSIz* tool (version 2.0) is employed with its default settings, utilizing the “-j” option for RIBOSUM scoring and the “-t” option for modeling transition/transversion ratios.

#### R-scape

*R-scape* employs simulated null alignments to assess the expected count of false positives in covariation analyses by employing a null hypothesis that maintains phylogenetic relationships while allowing columns in the alignments to evolve independently. This generates synthetic datasets that mirror the original sequences, facilitating the evaluation of the statistical significance of observed covariation patterns amidst random phylogenetically compatible variation. Through comparison of observed covariation scores with those expected under the null hypothesis, *R-scape* utilizes the G-test to distinguish genuine covariation signals from noise. *R-scape* version 2.0.4.a was run with the “s” flag to perform two independent statistical tests: one analyzing pairs included in the predicted structure and another examining all remaining possible pairs. The “lancaster” flag was also used to aggregate the *p*-values of base pairs in the same helices.

### Benchmarking

#### Predicting conserved RNAs in mitochondrial genomes

Mitochondrial genomes were analyzed using the *Enredo-Pecan-Ortheus* (EPO) consistency-based multiple sequence alignments of 46 eutherian mammals (63). These alignments were obtained from the ENSEMBL Compara release 104, which is publicly available online at (http://ftp.ensembl.org/pub/release-104/maf/ensembl-compara/multiple_alignments/46_mammals.epo) (64). The alignment data were first processed using *RNALalifold* and then with *SISSIz* version 2.0 and *R-scape* 2.0.4.a. We recorded the sensitivity of both *SISSIz* and *R-scape* at predefined thresholds.

*SISSIz* and *R-scape* predictions were considered true positives if they overlapped rRNA or tRNA annotations from GENCODE v.44 by at least 90% using *bedtools* (65). All other predictions were considered as false positives. Predictions were included for further analysis only if they demonstrated a Z-score of 0 or lower for *SISSIz*, or if they identified at least one significant base pair with an E-value less than 0.05 for *R-scape*. For helices, the log_10_ of the smallest E-value was retained.

Unless stated otherwise, the following intervals were selected to quantify accuracy at set thresholds for all benchmarks: *SISSIz* Z-scores from -11 to -1 in steps of 1; *R-scape* significant basepairs from 1 to 11 in steps of 1; *R-scape* helix log_10_(E-val) from -15 to 1 in increments of 2.

#### Simulated genome benchmarking of RFAM structures

Alignment blocks of 600 columns from the Ensembl compara alignments (release 104) with mean pairwise identity (MPI) between 85 and 95% were used to generate blocks of 600-nucleotide sequences that matched this MPI criteria, and then shuffled sequences from the Ensembl alignments using the *SISSIz* ’-s’ and ‘-t’ commands.

The complete alignments of 127 structural RNA families with 3D structures — excluding miRNAs and any alignments shorter than 50 nucleotides — were downloaded from Rfam release 14.9 (https://ftp.ebi.ac.uk/pub/databases/Rfam/14.9). Our algorithm, called *RfamConcealer*, was designed to randomly select an Rfam fasta file, from which subsets of 5, 10, or 20 sequences were randomly extracted. These subsets, characterized by MPI ranges of 50-65%, 65-80%, and 80-95%, were then integrated at position 300 of the shuffled EPO alignments to create hybrid blocks.

The hybrid blocks were then stripped of gaps and underwent multiple sequence alignment with *mafft-ginsi* (release 7.505) (66), thus reproducing the experimental conditions of a genomic screen and serving as a standardized control to assess the impact of sequence alignment on consensus RNA structure prediction tools. Subsequently, these alignments served as input for the *SISSIz* and *R-scape* tools. We calculated the mean positional index of each Rfam family within the various rows of the alignments. Using *bedtools* (65), we classified outputs as true positives if they were overlapped by 90% or more by the predicted inserted range within the hybrid alignment. All other predictions were deemed as false positives.

To evaluate the raw performance of our tools, we computed the area under the curve (AUC) of the receiver operating characteristic (ROC). The ROC curve is created by plotting sensitivity against the false positive rate (1-specificity) at various threshold values, helping to gauge the tool’s effectiveness in predicting RNA secondary structure. Following established practices (67), we also calculated sensitivity and positive predictive value (PPV). The F_1_ score, which is the harmonic mean of PPV and sensitivity, was then determined to further assess performance.

### Feature selection and machine learning

The simulated genomic benchmarking alignments were grouped into training and testing sets, with 80% of the alignments being allocated for training and 20% for testing. Alignments were scanned in both orientations with *RNALalifold*, generating over 29, 000 consensus secondary structures. The following features were selected for subsequent machine learning: *RNALalifold*’s consensus MFE score (thermodynamic stability); *RNALalifold*’s pseudo-energy (a heuristic measure of covariation); *SISSIz*’s Z-score (likelihood of structural conservation), the mean consensus MFE of *SISSIz* shuffled alignments (background thermodynamic stability), standard deviation of the background MFE (consistency of background alignment MFE); *R-scape*’s number of significant base pairs (a measure of covariation); *R-scape’*s minimum log_10_(E-value) (compounded covariation in helix); and the MPI.

A Generalized Linear Model (GLM) was implemented using the *glm* method from the *caret* package in R (68). 10-fold cross-validation was used to optimize model training, employing the *trainControl* function from the *caret* package. The *randomForest* method of the *caret* package was also used to train a random forest (RF) classifier with 10-fold cross-validation to optimize model performance and avoid overfitting. The model was tuned using mtry=5 with the best model selected based on the AUC of the ROC.

Feature importance was assessed using the iml package for permutation importance. We computed the drop in model performance (1 - AUC of the ROC curve) when each feature was randomly permuted. This permutation process was repeated 20 times for each feature to obtain mean importance scores and 95% confidence intervals. The machine learning models were evaluated in the context of different MPI and number of species ranges by dividing all alignments into distinct range groups. For each range group (MPI and the number of species) the GLM and RF were trained on all other range groups and applied to the group in question to evaluate the generalizability of each model.

#### ECSfinder

We designed *ECSfinder* as an opensource Java-bundled pipeline for the analysis of consensus RNA secondary structures from multiple genomic sequence alignments incorporating the optimal results presented herein by default. *ECSfinder*’s usage is dynamic and can be tuned via user-provided parameters. It requires a Multiple Alignment Format (MAF) file as input that is subjected to the *RNALalifold* tool from the Vienna RNA package (23) (version 2.4.16, available at (https://www.tbi.univie.ac.at/RNA) from which locally stable RNA secondary structures are extracted. This computation incorporates the RIBOSUM scoring metric and is constrained by user-defined parameters. In this study, we employed a maximum base pair span between 50 and 300 nucleotides (nt) and the exclusion of lone pairs.

To ensure the accuracy and relevance of the predicted secondary structures, both sense and antisense alignments were examined and compared against a background distribution created from dinucleotide-shuffled alignments using *SISSIz* (version 2.0). Alignments are subsequently processed with *R-scape* before extracting the relevant features used in the RF classifier. Default output is a list of alignment coordinates with associated prediction scores.

The *ECSfinder* pipeline and associated benchmarking scripts are available at https://github.com/TheRealSmithLab/ECSfinder.git.

## Results

### Detecting known RNA secondary structures in mitochondrial genomes

The mitochondrial genome is well-suited to assess the precision and efficacy of algorithms that predict functional noncoding RNA secondary structures. This compact genome is easy to align given its size and synteny, while being subject to robust selective pressures that retain functional structured RNAs across a wide range of species. It harbors well-annotated transfer RNAs (tRNAs) that are intercalated between messenger RNAs (mRNAs) and ribosomal RNAs (rRNAs); their position is essential for the processing of polycistronic mitochondrial RNAs via post-transcriptional endonuclease cleavage (61). We therefore utilized mitochondrial tRNAs and rRNAs as controls to empirically assess the performance of *SISSIz* and *R-scape* at identifying evolutionarily conserved RNA structures, as generated by *RNALalifold*.

Mitochondrial sequence alignments from eutherian mammals were first queried with *RNALalifold* to predict locally stable RNA secondary structures using maximal base pair distances of 100, 200, and 300 nt. These generated 820, 888, 970 consensus structures (many of which were partially overlapping), respectively, when both sense and antisense alignments were considered. *SISSIz* and *R-scape* were then applied, reporting a total of 436 and 230 conserved RNA structures, respectively, at ≤300 nt using the most liberal statistical threshold, i.e., Z-score ≤ -1 and ≥ 1 significant consensus base pairs (Fig. 1A). In *SISSIz* predictions, structures overlapping tRNAs and rRNAs usually had lower Z-scores (approaching -11) compared to those overlapping mRNA annotations (which were conservatively dismissed as non-structural), yet some more significant overlaps within mRNA were nonetheless detected (Fig. 1B). A similar trend was observed for structures with maximum base pair lengths of 100 and 200 (Supplementary Fig. 1). Although *R-scape* did not identify significant base pairs in predictions within mRNAs, it also did not report significant RNA base pair predictions in the mitochondrial tRNAs for isoleucine (MT-TI), glutamine (MT-TQ), or methionine (MT-TM). Interestingly, *RNALalifold* failed to produce consensus secondary structure predictions for a single mt-tRNA, the glutamic acid tRNA (MT-TE).

**Figure 1.**
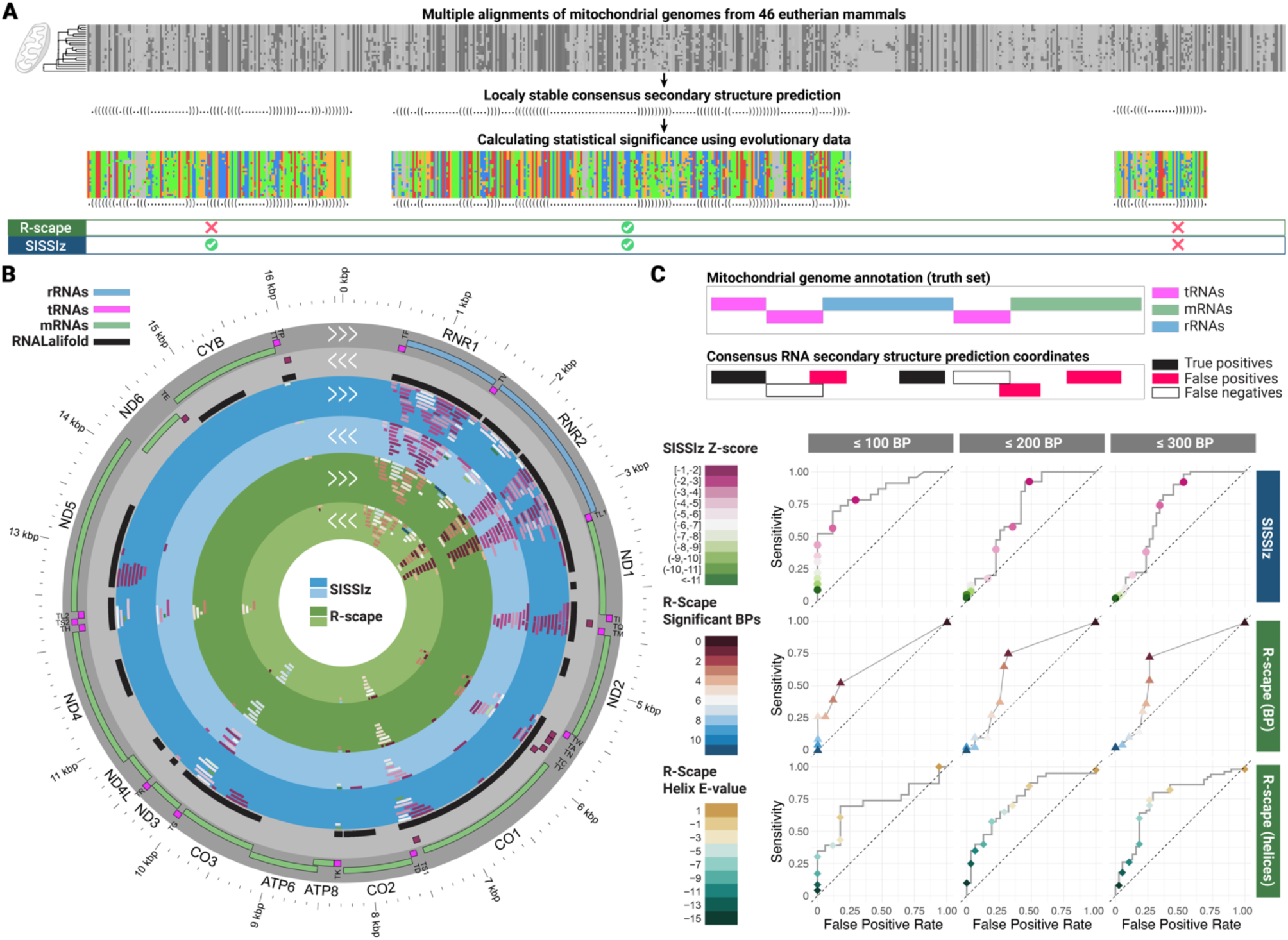
Predicting conserved ncRNA structures in mitochondrial genomes for maximum 300 base pairs in length. (A) Overview of benchmarking pipeline. Multiple genome sequence alignments are input into *RNALalifold* to predict locally stable secondary structures, which are then scored for evolutionary conservation by *SISSIz* and *R-scape*, then compared to reference gene annotations. (B) *Circos* (73) map of the human mitochondrial genome with the following tracks (from outward in): Gene annotation; RNA secondary structures predicted by *RNALalifold* (collapsed in black); ECS predictions by *SISSIz*, annotated for both positive (dark gray) and negative (light gray) strands; ECS predictions by *R-scape*. Triple arrows indicate the sense of the strand (positive or negative). (C) ROC curves comparing the performance of *SISSIz* and *R-scape* in identifying true positive RNA secondary structures across three different maximum base pairs lengths: ≤100 bp, ≤200 bp, and ≤300 bp.

Predictions with ≥90% overlap with rRNA or tRNA sequences were classified as true positives. Although some ECS predictions in mRNA and other noncoding sequences might represent conserved RNA structures (69–72), in the absence of experimental validation, all predictions other than mitochondrial tRNAs and rRNAs were considered false positives. Notably, *SISSIz* achieves higher sensitivity at a maximal threshold of - 1 compared to *R-scape* at minimum 1 significant base pair across all three conditions (100, 200, and 300 maximum base pairs) (Fig. 1C). However, this increase in sensitivity comes with a slightly higher false positive rate at equivalent true positive rates. Among the 436 structures identified by *SISSIz* with a Z-score ≤ -1, 214 were categorized as false positives, with 59% of these overlapping 5’ or 3’ untranslated regions. When comparing the AUC for the individual features, *SISSIz* outperformed *R-scape* in terms of the number of significant base pairs in all three windows, both performing best at base pair distance ≤100 nt with an AUC of 0.84 (*SISSIz*) versus 0.69 (*R-scape* base pairs) (Table 1), which is expected given that the average length of tRNAs is around 70 nucleotides.

**Table 1.**
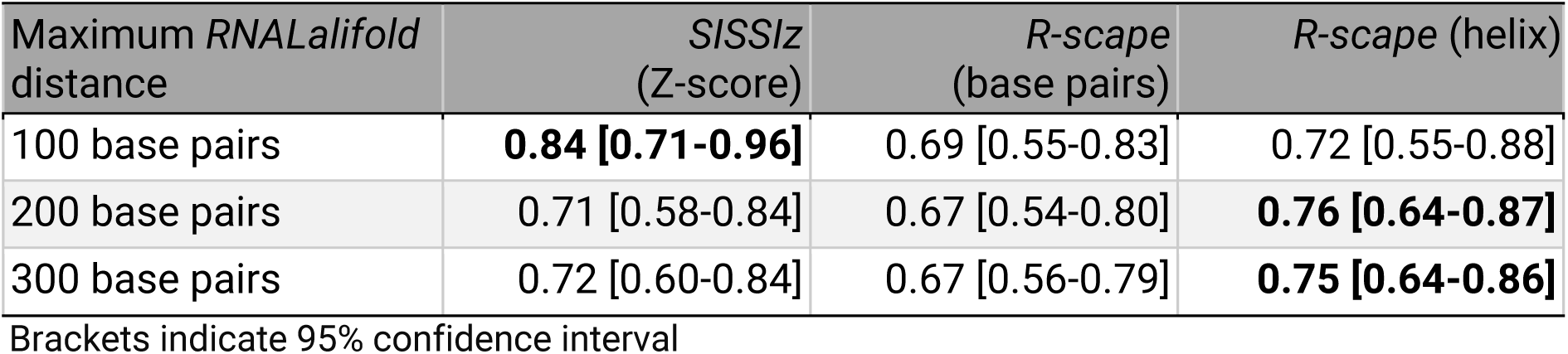
AUC of mitochondrial ncRNA ECS predictions.

Additionally, we applied a novel functionality of *R-scape* that enables the identification of significant RNA helices based on aggregating *p*-values of base pairs in proximity (39). The reported helix E-value, which accounts for multiple testing, multiplies the *p*-value of the helix by the total number of helices in the proposed structure. Thus, it estimates the expected number of false helices that would have such E-value or larger. Considering the minimum E-value of each predicted structure and taking the log_10_ of these values significantly increased the accuracy of predictions. The AUC is improved by 13.4% for the 200 base pairs condition, increasing from 0.67 when considering the number of base pairs to 0.76 when considering the lowest E-value in the predictions (Table 1). Interestingly, not all helices determined to be significant by *R-scape* contained ‘significant base pairs’, suggesting that helix scoring is a more accurate metric.

The parallels and differences in the outputs of both tools are exposed in two examples: one showcasing an ECS prediction identified by both tools, the mt-tRNA lysine (MT-TK; Fig. 2A); the other highlighting a *SISSIz*-specific prediction, the mt-tRNA glutamine (MT-TQ; Fig. 2B). Both secondary structures are accurately predicted by *RNALalifold* and reveal base-pair covariation in the alignments that are compatible with the inferred phylogeny, indicative of conservation of molecular function conveyed through higher-order structure.

**Figure 2.**
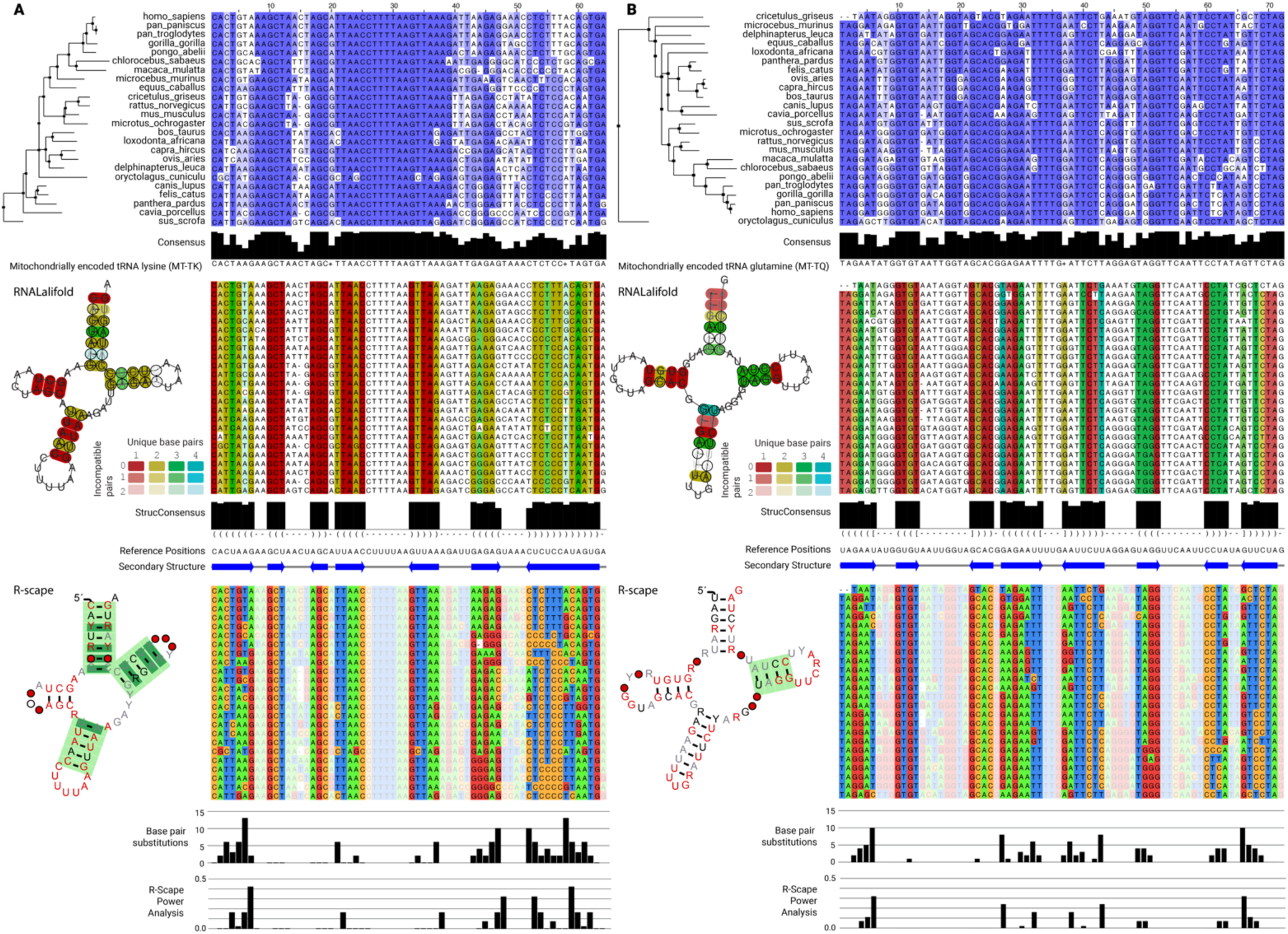
Visualization of two different predictions in the mitochondrial genome. (A) Mitochondrially encoded tRNA lysine. The top alignment displays percentage identity. The middle alignment shows the output from *RNALalifold*, with columns color-coded based on the number of unique base pairs within the predicted secondary structure. The reference sequence and structure from *RNALalifold* are also included. The bottom alignment is color-coded according to nucleotide type. Base pair substitutions and power analysis results are derived from the output of *R-scape*. Helices predicted to be significant by *R-scape* are highlighted in dark green (helix aggregated E-value <0.05) at the bottom panel of the figure. (B) Mitochondrially encoded tRNA glutamine, presented in the same format as (A). Alignments visualized with *Jalview* (75).

For MT-TK (Figure 2A), *RNALalifold* reports a consensus folding energy of -19.21 kcal/mol with *SISSIz* producing a Z-score of -10.44 (background mean of -1.02 kcal/mol, as detailed in Supplementary Fig. 2A). This examination uncovered one base pair that exhibits four distinct types of base pair interactions, alongside two base pairs demonstrating three unique types, and nine base pairs showcasing two unique types (Fig. 2A, middle panel). Additionally, we evaluated the alignment power using *R-scape*, which represents the fraction of base pairs expected to demonstrate significant covariation signals. *R-scape*’s two-set covariance test, which is more sensitive than its default one-set test when the structure is known *a priori*—as in the case with *RNALalifold*’s generated structure—contrasts the base pairs in the consensus structure against all other possible pairs. *R-scape* recognized eight significantly covarying base pairs (in dark green) out of the 20 proposed by *RNALalifold* despite it being a “low power alignment” (below 10%) according to (74). This number remains lower than the 20 base pairs identified by *RNALalifold*. Moreover, using the Lancaster aggregation method (39), 3 out of the 4 helices were determined to be significant (shown in light green in the lower panel of Fig. 2A).

For MT-TQ (Fig. 2B), *RNALalifold* reports a consensus folding energy of -15.80 kcal/mol against a mean background score of -2.28 kcal/mol, producing a *SISSIz* Z-score of -4.88 (detailed in Supplementary Fig. 2B). It identified one base pair with four unique types of interactions, six base pairs with three unique interactions, and three base pairs with two unique interactions. *R-scape* did not recognize any of these base pairs as significant despite some base pairs showing potential for significance based on power and substitutions. However, one helix containing no significant base pairs was identified as significant using Lancaster aggregation (E-value: 0.0216). Although of similar power to MT-TK, no significant base pair was observed in this structure. These results indicate that *SISSIz* can identify regions that are conserved at the secondary structure level that *R-scape* cannot when looking at the number of significant covarying base pairs.

### Predicting RNA structure conservation in simulated genomic alignments

Albeit practical for identifying conserved RNA secondary structures, the limited sequence diversity and evolutionary dynamics of polycistronic mitochondrial genomes differ from nuclear genomes. We therefore established a benchmarking workflow to emulate the properties of multiple genome sequence alignments, reflecting the experimental conditions typically observed in genome-wide comparative screens (see Methods and Fig. 3A). Randomly selected regions were extracted from Ensembl multiple sequence alignments of mammalian genomes and shuffled with SISSI. Next, we inserted stochastically sampled sequences from Rfam families with experimentally validated 3D structures within the shuffled genomic alignments. These hybrid blocks were then de-gapped and realigned using a sequence alignment tool prior to being queried with *RNALalifold*, *SISSIz*, and *R-scape*. We assessed the MPI of each Rfam family within the alignments and evaluated the concordance between the anticipated positions of Rfam families and the outputs provided by *SISSIz* and *R-scape*. Predictions that were at least 90% contained within the coordinates of the inserted Rfam sequences were classified as true positives. Conversely, predictions overlapping either extremity of these coordinates were categorized as false positives.

**Figure 3.**
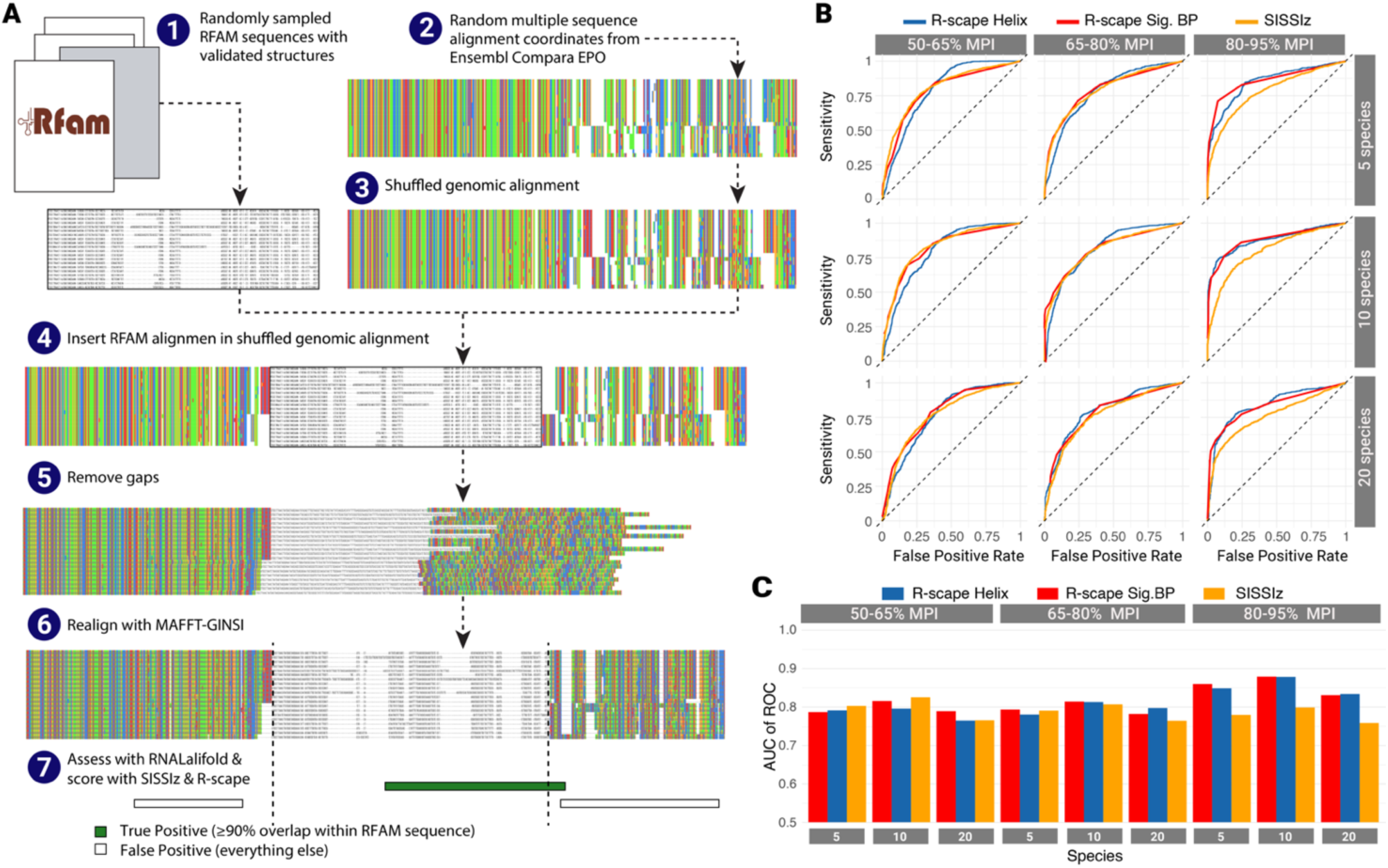
Simulated genomic ECS prediction benchmark. (A) Overview of benchmarking design and workflow. This analysis included Rfam families comprising 5, 10, and 20 species with MPIs ranging from 50 to 95%. (B) ROC curves for the different ECS prediction algorithms and scoring modalities against different MPI ranges and number of species. (C) Area under the curve values associated with (B).

We compared the Z-score from *SISSIz* against the minimum Lancaster E-value of each helix and the number of significant base pairs generated by *R-scape*. Overall, *SISSIz* and *R-scape* performed similarly when comparing the Z-score to the number of significant base pairs (Fig. 3B, C). The largest discrepancy between *SISSIz* and *R-scape* (significant base pairs) occurs in the 80-95% sequence identity range when analyzing 10 sequences with an AUC of 0.80 for *SISSIz* versus 0.88 for *R-scape* (Fig. 3B and Table 2). *SISSIz* performed best for RFAM alignments within the 50-65% MPI range with 10 species, achieving an AUC of 0.83, whereas *R-scape* performed best in default mode (significant base pairs) within the 80-95% MPI range with ≤ 10 species, with a maximal AUC of 0.88 (Table 2). However, both variants of *R-scape* presented similar AUC values for the various conditions. It should be noted that *R-scape*’s significant base pair scoring displays lower granularity in the ROC curves compared to *R-scape* helix and *SISSIz*, which reflects the limited range and discrete nature of the metric compared to the more diverse and continuous values of the other scoring metrics. Interestingly, all tools present slightly lower AUC values with 20 species, suggesting that sequence alignment may obfuscate informative covariation signals.

**Table 2.** AUC of *SISSIz* and *R-scape* across MPI ranges and number of species in simulated genomic alignments.

| MPI | Number of species | *SISSlz* (Z-score) | *R-scape* (base pairs) | *R-scape* (helix) |
| --- | --- | --- | --- | --- |
| 50-65% | 5 | **0.804 [0.791-0.816]** | 0.788 [0.775-0.800] | 0.792 [0.780-0.804] |
|  | 10 | **0.826 [0.812-0.840]** | 0.816 [0.802-0.831] | 0.797 [0.782-0.811] |
|  | 20 | 0.766 [0.746-0.786] | **0.790 [0.772-0.808]** | 0.765 [0.745-0.785] |
| 65-80% | 5 | 0.791 [0.778-0.804] | **0.794 [0.782-0.807]** | 0.781 [0.768-0.794] |
|  | 10 | 0.807 [0.793-0.822] | **0.815 [0.801-0.829]** | 0.814 [0.800-0.828] |
|  | 20 | 0.765 [0.745-0.785] | 0.783 [0.764-0.801] | **0.798 [0.779-0.817]** |
| 80-95% | 5 | 0.780 [0.763-0.798] | **0.861 [0.846-0.875]** | 0.849 [0.834-0.864] |
|  | 10 | 0.800 [0.778-0.821] | **0.880 [0.863-0.896]** | 0.879 [0.862-0.897] |
|  | 20 | 0.759 [0.740-0.779] | 0.832 [0.816-0.847] | **0.835 [0.818-0.851]** |

Brackets indicate 95% confidence interval

An important factor to consider when performing genome-wide screens for ECSs is the orientation of predictions. It is common that ECSs are predicted on both strands (as can be observed in Fig. 1B) consistent with base-pairing complementarity, with the exception of G:U base pairs. We found that sequence properties and scoring metrics between (+) and (–) predictions were largely overlapping in our benchmark (see below), rendering it difficult to determine in which orientation the RNA structure is located. This prompted us to mine the various features of ECS predictions from multiple alignments to evaluate their discriminative potential.

### Improving ECS predictions with machine learning classifiers

We next explored if the subtle yet distinctive differences between *SISSIz* and *R-scape* predictions across a range of conservation, stability and covariation characteristics could be sufficient to improve strand-specific ECS prediction through classical machine learning (ML) strategies. We selected features derived from the alignment (MPI), consensus structure prediction with *RNALalifold* (MFE and pseudo-energy, a thermodynamic proxy of base-pair covariation), *SISSIz* metrics (Z-score, median *RNAalifold* MFE of the background sampling, and the standard deviation of the latter, a measure of the background’s consistency), and *R-scape* metrics (both the number of significant base-pairs and the minimum log_10_(E-value) of helix) (Fig. 4A).

**Figure 4.**
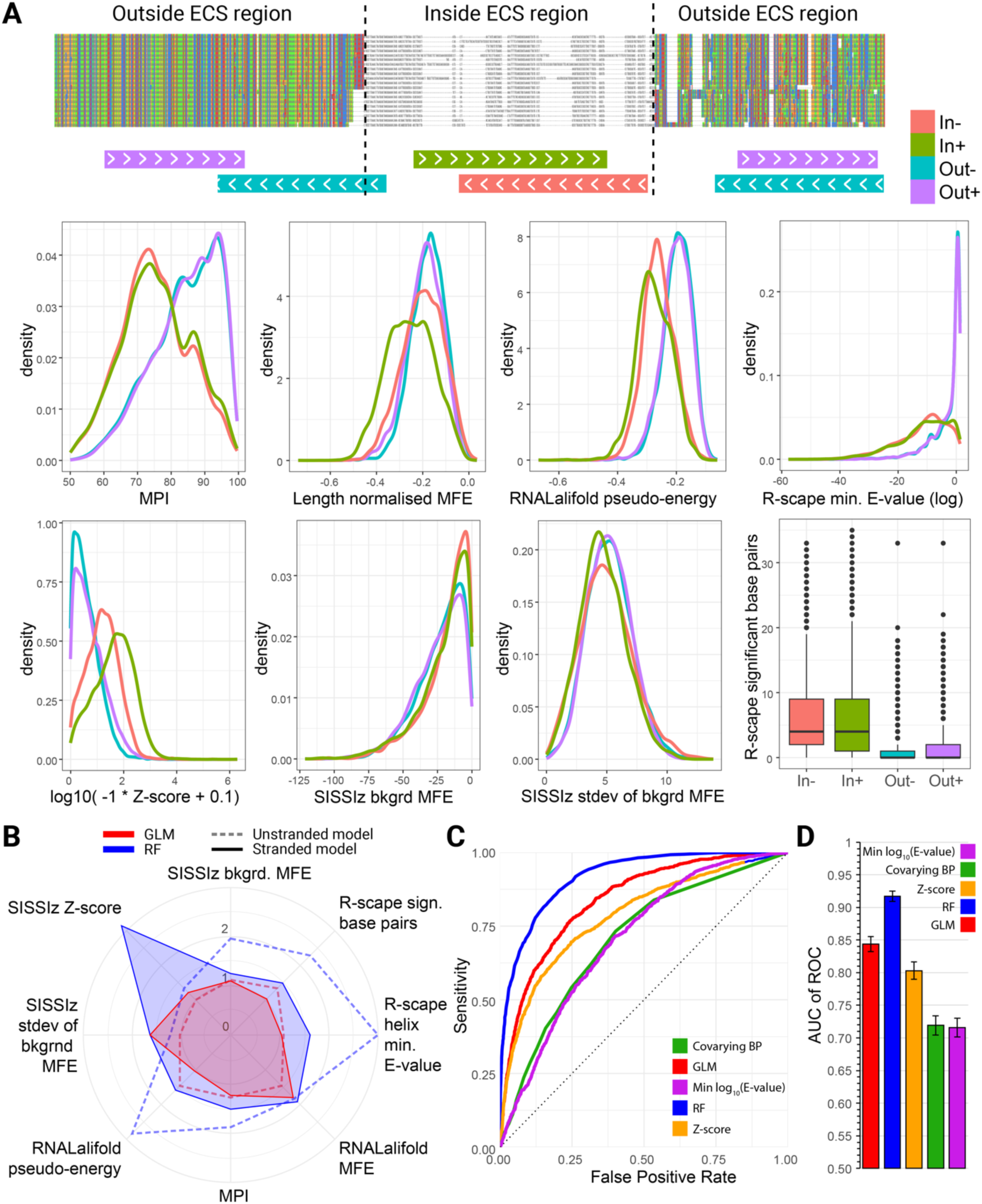
Feature selection and performance of strand-aware ECS machine learning classifiers. (A) Summary of prediction classification (top) and distribution of selected features used for ML training. (B) Radar plot showing the relative importance of predictive features across both ML models, with and without considering strandedness. Values indicate how each model’s performance (measured as 1 - AUC) decreases when the respective feature is randomly permuted (mean of 20 permutations displayed). Features with higher importance contribute more significantly to the predictive power of each model. (C) ROC curves comparing the performance of strand-aware ECS prediction methods across all MPI and number of species ranges. ROC values were calculated on a distinct validation set (20% withheld data) following training using a leave-one-out 10-fold cross validation paradigm. (D) Area under the curve values of (C) with error bars indicating 95% confidence intervals.

These features were used to train a generalised linear model (GLM) and a random forest classifier (RF) to improve the identification of ECSs from *RNALalifold* predictions (see Methods). Two modalities were tested: (i) predictions were labelled as true positives if they were within the RFAM sequence coordinates of the alignment or false positives otherwise, and (ii) similarly, but considering only sense (+) orientated predictions within the RFAM sequence coordinates as true positives. To pinpoint the most critical features, we examined the relative importance of all features for both models, with and without consideration of strandedness (Fig. 4B). Both unstranded classifiers (dashed lines) exposed the importance of *RNALalifold*’s pseudo-energy as a top feature to identify genomic regions harbouring ECSs. However, the GLM model assigns relatively similar importance to *RNALalifold*’s MFE and *R-scape*’s significant base pairs features. The standard deviation of background MFE and Z-score metrics from *SISSIz* are negligeable contributors to the unstranded ML classifier models. Interestingly, the E-value metric is almost 3-fold more important for the RF than the GLM.

In contrast to the unstranded classifier, the *SISSIz* Z-score is the predominant discriminator in the stranded RF. This is justified by the Z-score distributions depicted in Fig. 4A, where a clear shift is observed for (+) versus (–) stranded predictions within the RFAM coordinates. The *RNALalifold* consensus MFE is also an important discriminative feature for both stranded ML models (also evidenced by a shift in kernel density distribution), suggesting that thermodynamic features are important to consider when considering strandedness. Interestingly, the standard deviation of the background MFE generated by *SISSIz* is the second and third most important feature for the stranded GLM and RF classifiers, respectively, despite contributing little to the discriminative ability of the unstranded models. This suggests that combining *SISSIz*’s Z-score with a measure of consistency of its background modeling, such as the standard deviation of the background MFE, can identify false positive predictions with more ease.

Although the GLM classifiers include a more balanced distribution of features in their scoring, the RF model provided the best classification accuracy, substantially outperforming all other ECS prediction tools (Fig. 4C, D and Table 3). The resilience of the RF model is evidenced by inspecting the performance of the different ML and non-ML ECS prediction methods across all surveyed MPI and number of species ranges, where no significant differences are observed (Supplementary Fig. 3). These findings, which combine data across different MPI ranges and species counts, underscore the effectiveness of ML models in achieving higher accuracy and better overall performance in the functional characterization of non-coding RNA structures compared to traditional, non-ML methods.

**Table 3.** Comparative performance of the ECS prediction and classification models.

| Threshold | Model | Cut-off | Sensitivity | Specificity | Accuracy | F1-score |
| --- | --- | --- | --- | --- | --- | --- |
| 1% FPR | GLM | 0.87 | 0.182 | 0.990 | 0.791 | 0.3 |
|  | Random Forest | 0.75 | **0.354** | 0.990 | **0.834** | **0.511** |
|  | *S/SS/z* | -12 | 0.133 | 0.990 | 0.562 | 0.233 |
|  | *R-scape* BP | 19.5 | 0.036 | 0.990 | 0.513 | 0.068 |
|  | *R-scape* Helix | -32.32 | 0.030 | 0.990 | 0.509 | 0.055 |
| 5% FPR | GLM | 0.57 | 0.440 | 0.950 | 0.824 | 0.552 |
|  | Random Forest | 0.51 | **0.615** | 0.950 | **0.869** | **0.697** |
|  | *S/SS/z* | -7 | 0.367 | 0.950 | 0.659 | 0.512 |
|  | *R-scape* BP | 11.5 | 0.152 | 0.950 | 0.551 | 0.248 |
|  | *R-scape* Helix | -21.99 | 0.138 | 0.950 | 0.544 | 0.226 |
| Maximum accuracy | GLM | 0.45 | 0.530 | 0.917 | 0.828 | 0.599 |
|  | Random Forest | 0.53 | 0.600 | **0.957** | **0.869** | 0.692 |
|  | *S/SS/z* | -4 | 0.709 | 0.764 | 0.736 | **0.702** |
|  | *R-scape* BP | 2 | **0.757** | 0.588 | 0.672 | 0.654 |
|  | *R-scape* Helix | -4.83 | 0.725 | 0.593 | 0.659 | 0.616 |
| Maximum F1-score | GLM | 0.31 | 0.688 | 0.848 | 0.809 | 0.758 |
|  | Random Forest | 0.38 | 0.789 | **0.889** | **0.864** | **0.864** |
|  | *S/SS/z* | -3 | 0.747 | 0.715 | 0.731 | 0.704 |
|  | *R-scape* BP | 1 | 0.854 | 0.454 | 0.654 | 0.659 |
|  | *R-scape* Helix | -0.75 | **0.890** | 0.386 | 0.642 | 0.631 |
| Youden Index | GLM | 0.26 | 0.740 | 0.806 | 0.789 | 0.629 |
|  | Random Forest | 0.28 | **0.869** | **0.825** | **0.838** | **0.722** |
|  | *S/SS/z* | -4 | 0.709 | 0.764 | 0.736 | 0.702 |
|  | *R-scape* BP | 2 | 0.757 | 0.588 | 0.672 | 0.654 |
|  | *R-scape* Helix | -4.83 | 0.725 | 0.593 | 0.659 | 0.616 |

## Discussion

In this study, we set out to compare two leading RNA secondary structure prediction tools—*SISSIz* and *R-scape*—which use different approaches to assess RNA structure conservation. *SISSIz* relies on thermodynamic modeling and covariation analysis across entire RNA structures, whereas *R-scape* focuses on detecting statistically significant base pair covariation. These complementary methods were tested using two distinct benchmarks: well-annotated mitochondrial RNA structures and experimentally validated RNA structure alignments embedded within simulated genome alignments. Our evaluation revealed that while both tools performed reasonably well in isolation, each had limitations affecting their ability to reliably predict evolutionarily conserved RNA secondary structures.

Overall, our results suggest that both methods perform similarly. *SISSIz* offers higher sensitivity by focusing on global thermodynamic stability, though this comes at the cost of more false positives. In contrast, *R-scape* provides greater specificity through covariation analysis but often misses broader structural conservation. Recognizing the distinct strengths and weaknesses of each tool, we developed *ECSfinder* to harmonize these methods. Rather than relying solely on one tool, *ECSfinder* integrates the complementary features of *SISSIz* and *R-scape* through machine learning. Specifically, *ECSfinder* combined thermodynamic stability and background modeling measures from *SISSIz*, covariation metrics from *R-scape*, and additional features such as *RNAalifold*’s pseudo-energy score and sequence conservation. By training a random forest model on these combined metrics, *ECSfinder* balances sensitivity and specificity, offering a more robust and reliable solution for RNA secondary structure prediction.

The comparative performance of *SISSIz* and *R-scape* was first evaluated using mitochondrial genomes, which serve as an ideal model due to their well-characterized RNA structures and evolutionary conservation. *SISSIz* demonstrated strong sensitivity in detecting RNA secondary structures, identifying a broad range of conserved structures with low Z-scores, particularly within mitochondrial tRNAs. However, this came with a trade-off: *SISSIz* generated a higher rate of false positives than *R-scape*. The conservative nature of our analysis should be emphasized, as any prediction that did not overlap a tRNA or rRNA by 90% was considered a false positive, including structures predicted within mRNAs that may in fact harbour conserved secondary structures (76). *R-scape*, focusing on significant base pair covariation, showed better specificity but failed to detect certain known conserved structures, such as tRNA helices, where covariation patterns were not pronounced. *R-scape*’s helix scoring noticeably alleviates this. The simulated genomic alignment benchmark demonstrated that both *SISSIz* and *R-scape* perform similarly across the surveyed conditions, with slightly better performance for *R-scape* in the 80-95% MPI range. These findings suggested that neither method alone was sufficient to accurately predict all evolutionarily conserved RNA structures, prompting the need for a new integrated approach.

*ECSfinder* was developed to address the complementary features of *SISSIz* and *R-scape*, combining *SISSIz*’s thermodynamic predictions and *R-scape*’s covariation signals into a unified model. Given the small size of the training sets and the complexity of deep learning models, we opted for machine learning methods that enhance interpretability, helping us understand how different predictors influence ECS detection. The feature importance analysis (Fig. 4) revealed that covariation signals were the most critical predictors of conserved RNA structures, highlighting the role of compensatory mutations in maintaining RNA structure across species. However, thermodynamic stability and the *SISSIz* Z-score also emerged as important for strand-specific ECS predictions, emphasizing the relevance of considering G:U ‘wobble’ base pairs in these models.

The role of pseudo-energy and MFE in feature importance is particularly intriguing, given the ongoing debate in the field about whether MFE should be considered when evaluating RNA for evolutionary conserved structure. It has been argued that when identifying RNAs with evolutionarily conserved structures, pure covariation measures like MI or the G-test are more reliable, as folding free energies are often inadequate for detecting highly structured RNAs—especially in longer sequences where free energy decreases linearly with sequence length (33, 77). However, our findings suggest that incorporating the physical properties of RNA, such as MFE, plays a meaningful role in accurately predicting conserved structures.

The development of *ECSfinder* opens exciting opportunities for further refinement and application. In its current form, *ECSfinder* is designed to work within comparative genomic alignments, but future versions could integrate additional experimental data, such as SHAPE-MaP or chemical probing, to further enhance prediction accuracy. Additionally, *ECSfinder* could be expanded to include more sequence properties, prediction metrics, alternative consensus structure predictions and more sequence alignment tools.

Ultimately, we improved overall prediction accuracy beyond what either tool could achieve individually. The enhanced AUC scores across benchmarks confirmed that the random forest classifier significantly outperforms the standalone use of *SISSIz* or *R-scape*, particularly in predicting the orientation (strandedness) of RNA structures. *ECSfinder* effectively captures both global and localized structural conservation, significantly improving sensitivity and specificity.

This study not only addresses the limitations of existing tools but also sets the stage for future advancements in RNA biology and computational genomics. *ECSfinder* can be used in large-scale comparative genomics studies to identify conserved RNA structures, including long non-coding RNAs (lncRNAs), which are increasingly recognized as key regulators of cellular function. By improving upon existing prediction methods, *ECSfinder* facilitates the discovery of novel functional RNAs and helps clarify the role of conserved secondary structures in gene regulation, development, and disease.

## Acknowledgements

We extend our thanks to Shaun Lovejoy for his meticulous reading of the manuscript and his valuable edits. Additionally, we thank Alexis Nolin Lapalme for insightful scientific discussions.

## Supplementary figures

**Supplementary Figure 1.**
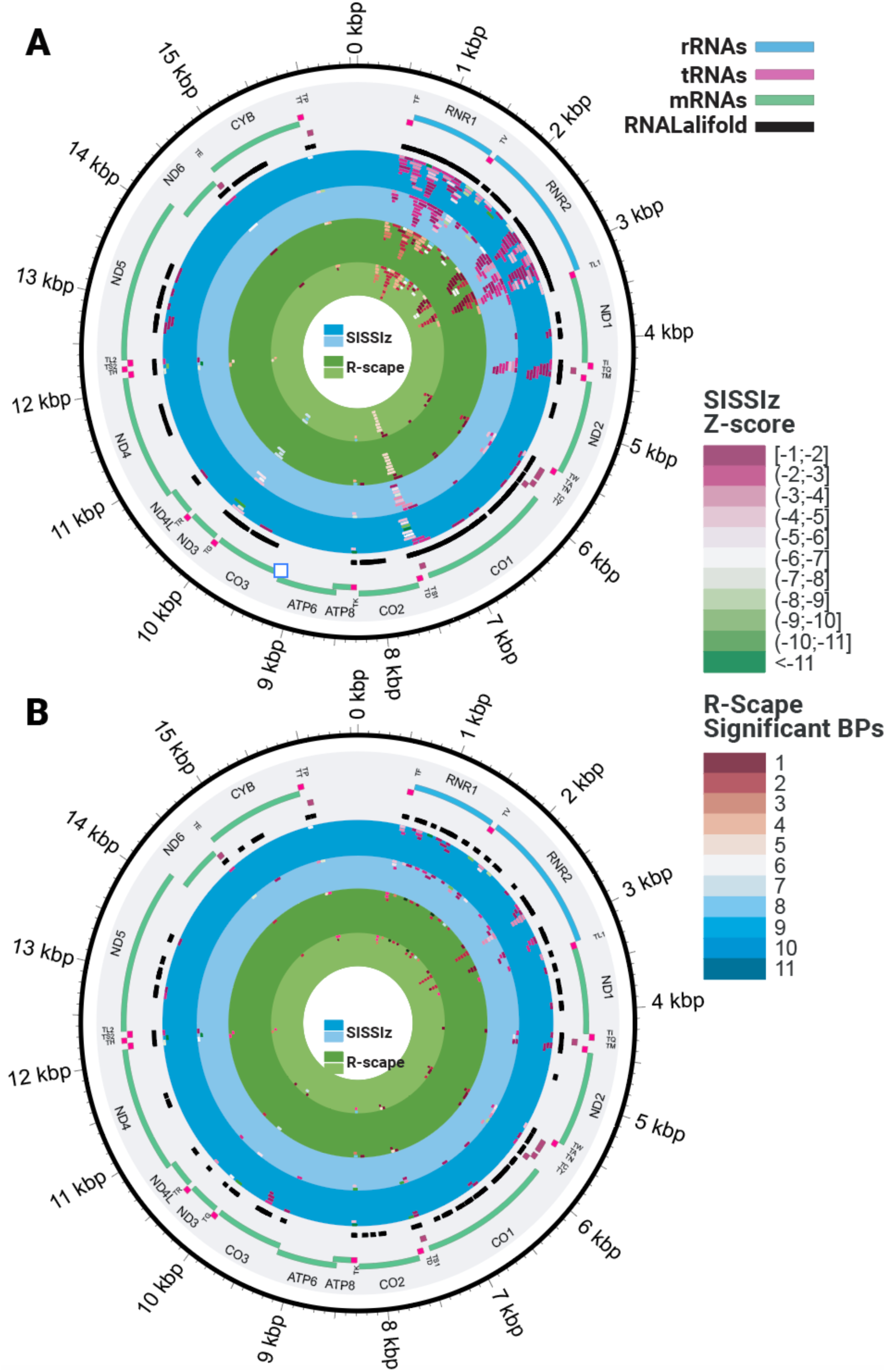
Benchmarking ncRNA discovery using the mitochondrial genome. (A) Analysis with a maximum of 100 nucleotides per base pair. (B) Analysis extended to a maximum of 200 nucleotides per base pair.

**Supplementary Figure 2.**
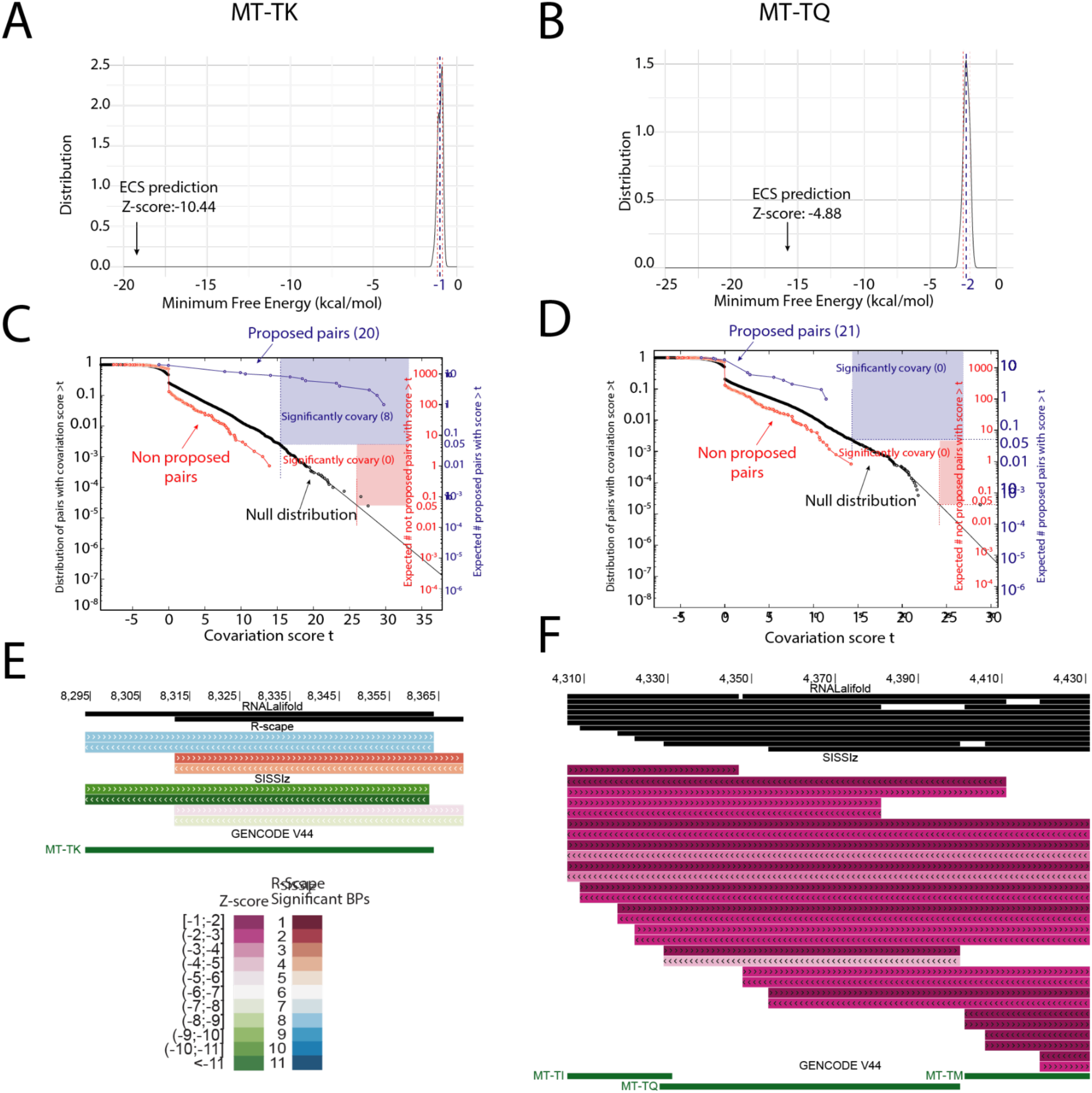
Comparative analysis of two mitochondrial tRNA predictions using SISSIz and R-scape. (A) Output from SISSIz for the mitochondrially encoded tRNA lysine (MT-TK), including the null background distribution and its mean MFE alongside the MFE of the consensus predicted structure. (B) Similar output from SISSIz for the mitochondrially encoded tRNA glutamine (MT-TQ). (C) Survival curve generated by R-scape for the prediction of MT-TK. The curve assesses statistical significance of base pairs within the predicted structure using the -s option: independent statistical tests are performed for the set of base pairs in the given structure (blue) and for the rest of the pairs (red). (D) Survival curve for the prediction of MT-TQ, displayed in the same manner as in (C). (E) UCSC Genome Browser track output showing the output predictions of SISSIz and R-scape overlapping the MT-TK. (F) UCSC Genome Browser track output for the MT-TQ.

**Supplementary Figure 3.**
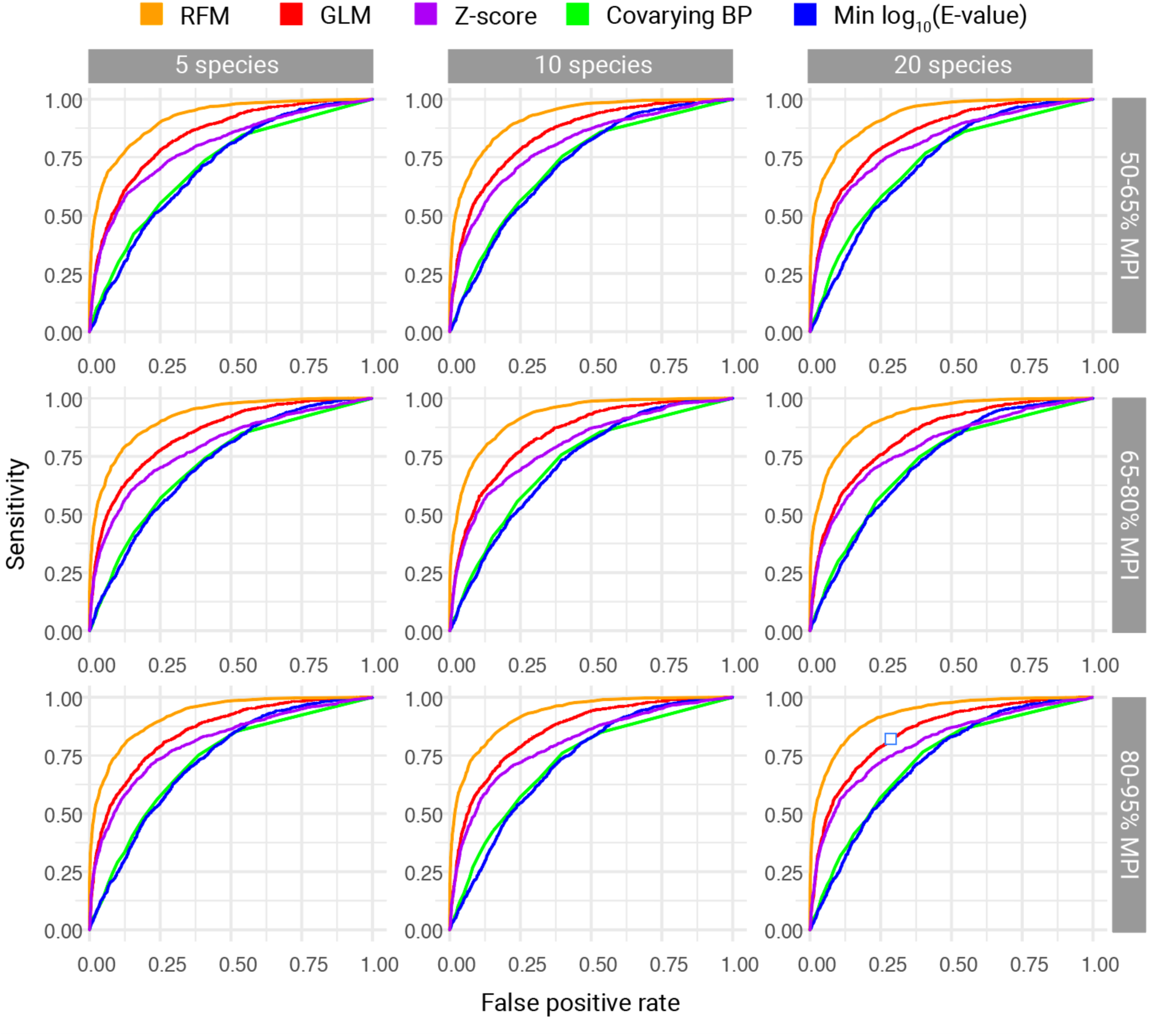
ROC curves for ML and non-ML ECS prediction methods in function of different sequence conservation ranges and number of species. Each plot represents the outcome of ‘leave one out’ training, where the evaluated MPI and number of species range was omitted from the respective training set.

